# Detecting presence of mutational signatures in cancer with confidence

**DOI:** 10.1101/132597

**Authors:** Xiaoqing Huang, Damian Wojtowicz, Teresa M. Przytycka

**Affiliations:** National Center of Biotechnology Information, National Library of Medicine, NIH, Bethesda MD 20894, USA

## Abstract

Cancers arise as the result of somatically acquired changes in the DNA of cancer cells. However, in addition to the mutations that confer a growth advantage, cancer genomes accumulate a large number of somatic mutations resulting from normal DNA damage and repair processes as well as mutations triggered by carcinogenic exposures or cancer related aberrations of DNA maintenance machinery. These mutagenic processes often produce characteristic mutational patterns called mutational signatures. Decomposition of cancer’s mutation catalog into mutations consistent with such signatures can provide valuable information about cancer etiology. However, the results from different decomposition methods are not always consistent. Hence, one needs to not only be able to decompose a patient’s mutational profile into signatures but also to establish the accuracy of such decomposition. We proposed two complementary ways of measuring confidence and stability of decomposition results and applied them to analyze mutational signatures in breast cancer genomes. We identified very stable and highly unstable signatures, as well as signatures that have been missed altogether. We also provided additional support for the novel signatures. Our results emphasize the importance of assessing the confidence and stability of inferred signature contributions. All tools developed in this paper have been implemented in an R package, called *SignatureEstimation*, which is available from https://www.ncbi.nlm.nih.gov/CBBresearch/Przytycka/index.cgi#signatureestimation.

## 1 Introduction

Knowledge of elementary mutational processes underlying cancer cells is essential for understanding the etiology of cancer and its progression. The ever-growing amount of sequencing data allows for the analysis of cancer genomes not only from the perspective of most frequently mutated genes, but also from the perspective of broader mutational patterns. It is increasingly recognized that there is a whole spectrum of mutational processes that contribute to the mutation landscape of cancers. Cancer genomes harbor, in addition to cancer driving mutations, other somatic mutations acquired during the normal cell cycle as well as these triggered by carcinogenic exposures such as tobacco smoking, ultraviolet light, replication stress, or by cancer related aberrations of DNA maintenance machinery such as mismatch repair and other factors. Each of these processes often leads to distinctive pattern of mutations – the so-called mutational signature. Computational methods developed to uncover such signatures from catalogs of somatic mutations [1–6], including the classical nonnegative matrix factorization (*NMF*) approach, build on the assumption that the mutations observed in a cancer genome are a result of several mutational processes and that various genomes might have different exposure to each of the contributing mutagens. Analyses of cancer genomes from the perspective of mutational signatures have been very informative. In particular, the APOBEC mutational signature being a footprint of activity of the APOBEC family of cytidine deaminases, is a key factor in many human cancers [7–9]. APOBEC activity has been proposed to be the direct cause of some cancer driving mutations [10–12]. As another example, a recent analysis of mutational signatures in genomes of tobacco smoking-associated cancers provided a new understanding of how smoking increases cancer risk [13].

Given the growing interest in studies of mutational signatures in cancers and the need for a better understanding of the relation between detected signatures and biological causes, it is important to confidently assign known signatures to patients and assess patients’ exposure to each of these signatures. As a step in this direction, Rosenthal *et al.* recently developed an approach, called *deconstructSigs* (*dS*), that given a patient’s catalog of mutations and a set of predefined mutational signatures determines a linear combination of the signatures that best reconstructs the patient’s mutational profile [14]. However, we observed that this decomposition is not always in an agreement with the signature contributions provided by the nonnegative matrix factorization approach. Importantly, while deconstructSigs is based on a heuristic, the decomposition problem can be solved using a quadratic programming (*QP*) approach [15] that rapidly converges to the optimal solution. We observed that the optimal decomposition is sometimes strikingly different from the decomposition provided by either of the two other approaches in terms of signature presence or inferred contribution of signatures in tumor samples. This prompted us to address the confidence and stability of the decomposition problem. In particular, our approach allows us to assess if a cancer genome was exposed to a given set of mutational signatures and to quantify the confidence in the estimated contribution of each mutational signature.

There are two complementary aspects related to evaluating the confidence and stability of a decomposition of mutational catalog into mutational signatures. The first is to measure decomposition variability and accuracy when the patient’s mutational catalog is perturbed, as mutational catalogs might be noisy and incomplete. In addition, mutational processes act in a stochastic way. Given that quadratic programming can quickly compute optimal solutions for the original and perturbed data, this question can be answered using a bootstrap analysis. The second and complementary perspective arises from considering equivalent approximate solutions. Note that computationally derived definitions of signatures cannot be assumed to be perfect as they heavily depend on the data utilized for signature discovery and the details of a signature deciphering method. Therefore, different linear combinations of signatures might equally well approximate a mutational profile observed in a given patient. If quantitatively different suboptimal solutions exist in close proximity to the optimal solution, our confidence in the biological relevance of the optimal decomposition is reduced. To address this concern, we used a simulated annealing based method to randomly explore alternative decompositions whose error is close to the optimal one.

We applied both approaches to the whole-genome dataset of somatic mutations from 560 breast cancer (BRCA) patients [16, 17]. We identified very stable signatures (e.g. APOBEC related signatures 2 and 13) and highly unstable signatures (such as signatures 3, 5 and 8). We found that unstable signatures can be decomposed into other signatures with relatively small error, which explains the lack of stability of these signatures. Next we re-analyzed the BRCA data and found signatures whose presence in breast cancer have been missed by previous analyses. For the missed signatures, in addition to statistical validation, we also evaluated their association with genomic features that provided additional support for these signatures. Our results emphasize the importance of assessing the confidence and stability of inferred signature contributions for the interoperability of the decomposition results.

## 2 Materials and Methods

### Decomposition of mutational profiles into predefined signatures

Given a mutational catalog of a cancer genome (a set of somatic mutations) we strive to identify operative mutagenic processes and to quantify genome’s exposure to each of them. The imprint of a particular mutational process, referred to as its signature, is defined by the relative frequency of all types of nucleotide substitutions typically within the context of specific flanking residues. Here, we utilized mutational signatures that have been already discovered [2]. The goal is to approximate a genome mutational profile (i.e. observed mutation frequencies) by a linear combination of the signatures. Each signature coefficient of the linear combination is interpreted as genome exposure (i.e. a fraction of mutations caused in a genome) to a mutagenic process represented by the respective signature.

Formally, let *M* (*K × G*) be a matrix containing observed mutational profiles from *G* samples (e.g. 560 breast cancer patients), where each profile contains frequencies of *K* mutation types (e.g. 96 substitution types in the trinucleotide context) computed from patient’s catalog of mutations; and let *P* (*K × N*) be a matrix of *N* predefined mutational signatures (e.g. 30 COSMIC signatures) defining probabilities of generating each of *K* mutation types by separate mutational processes. The objective is to find a nonnegative exposure matrix *E* (*N × G*) that contains for each of *G* samples its exposure to each of *N* signatures by minimizing a Frobenius norm: 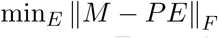; the value of the objective function represents the error of the inferred decomposition. For each patient *g* (*g* = 1,…,*G*), the objective function can be written as:

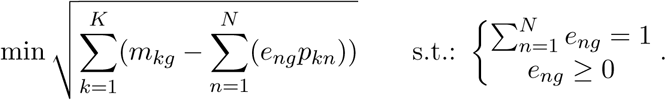

There are many approaches to solve such a minimization problem. In this paper, we implemented two methods based on quadratic programming (*QP*) and simulated annealing (*SA*); for details see Supplementary Materials. We used two publicly available R packages (https://cran.r-project.org): *quadprog* and *GenSA* for QP and SA, respectively. Both algorithms can find the optimal solutions. QP is extremely fast and stable, but it requires a predefined signature matrix *P* to be full column rank. SA can be widely used on a not-well-defined signature matrix, but it is slower than QP in converging to the optimal solution. Importantly, SA can be used to explore the landscape of suboptimal solutions that are close to the optimal decomposition.

### Confidence and stability of signature contributions

Biological data are noisy and the best way to quantify this uncertainty is to replicate the measurements, but that is not always possible. Instead, we used the bootstrap to determine how estimates of signature contribution were distributed and to answer questions about their confidence and stability. We perturbed the original mutational catalog of each patient 1000 times by randomized re-sampling with replacement and we estimated signature contributions in each bootstrap sample using QP method. Based on the distribution of signature contributions, one can estimate their bootstrap confidence intervals and the empirical probability that a signature contribution is above a specific threshold. To assess the stability (bias) of contributions for different signatures, i.e. to test how much the contributions of bootstrap experiments vary from the original contributions, we computed the mean squared error (*MSE*) of the difference between the bootstrap estimates and the optimal contributions in the original data.

Apart of decomposition stability with respect to the input data, we tested the stability of the optimal solution itself to see if there are any hidden dependencies between signatures. We sampled the space of suboptimal decompositions, i.e. approximate decompositions having slightly higher error than the optimal solution, using the simulated annealing approach – the same GenSA implementation we used to find the optimal solutions but with modified stopping rule allowing to report suboptimal solutions when a given value of objective function (decomposition error) is reached. The decomposition error threshold was set to be higher by factor of 1%, 3% or 5% relative to the error of the optimal solution. The calculations were repeated 1000 times for each patient to randomly sample the space of suboptimal decompositions with a given increase of decomposition error. The obtained distributions of signature contributions were then used, similarly to the bootstrap analysis, to assess confidence and stability of signature contributions.

### Genomic features of signatures

Morganella *et al.* [17] showed that breast cancer mutational signatures exhibit distinct relationships with genomic features related to transcription, replication and chromatin organization. Those analyses indicate that, in addition to sequence context captured by the nucleotides flanking each mutation, more general genomic context also matters. In particular, if mutational profile of a patient is shaped by several mutational processes then mutations from the contributing processes are not shuffled randomly over the genome. Instead, mutations from each operating process are often clustered together. Maximal stretches of adjacent substitutions in a sample generated by the same mutational process on the same reference allele are called processive groups. Our analysis of breast cancer data (see Results) has identified previously missed signatures, thus, in addition to statistical analysis we tested these signatures for expected genomic features, following procedures of Morganella *et al.* as summarized below (for details see [17]).

Each base substitution was associated with a mutational signature having the highest *a posteriori* probability of generating this mutation; the *a posteriori* probability was computed based on signature exposure levels inferred by QP method. Mutations (in the pyrimidine context) within protein coding genes (Ensembl release 60) were classified based on whether they are located on the transcribed/non-coding strand or the non-transcribed/coding strand. Transcriptional strand bias was computed for each signature separately as a ratio of the number of mutations on the transcribed strand to the total number of mutations in all samples. For processive groups, we counted the number of groups of different lengths (the number of maximal successive mutations) for each signature separately. To assess significance of the observed group counts, we compared them with the numbers of processive groups in 100 randomized dataset, where the order of mutations was shuffled with respect to the original data.

### Data sets

Whole-genome dataset of somatic mutations from 560 breast cancer (BRCA) patients [16, 17] was downloaded from ICGC Data Portal, https://dcc.icgc.org/releases/current/Projects/BRCA-EU (release 23). We classified all somatic substitutions into 96 mutation types in the trinucleotide context (6 substitutions from pyrimidine base pair times 4*4 nucleotide types at both 5’ and 3’ sides of substitution). For each patient we computed its mutational profile from the patient’s catalog of mutations, i.e. the number of mutations of each type. All patient’s mutational profiles were normalized by the total number of mutations that each contained.

The patterns of 30 known and validated mutational signatures were retrieved from the COSMIC website, http://cancer.sanger.ac.uk/cosmic/signatures (release 80). This set of signatures was deciphered from dozens distinct types of human cancer, although not all signatures are present in every cancer genome [1–5, 16]. The most recent analysis of the breast cancer whole-genome sequences by Nik-Zainal *et al.* [16] revealed 12 signatures found in breast cancer patients – signatures 1, 2, 3, 5, 6, 8, 13, 17, 18, 20, 26 and 30, and it provided exposures of each breast cancer patient to each signature as estimated by the NMF-based Mutational Signatures Framework [2]. We refer to this decomposition as *NMF decomposition*. We also applied the deconstructSigs approach to this dataset using 12 known breast cancer signatures to infer their exposure contributions [14]; none signature was discarded to minimize decomposition error. We refer to these estimates as *dS decomposition*.

## 3 Results

### 3.1 Differences in signature exposures inferred by different decomposition methods

The most commonly used tool for discovering mutational signatures from catalog of mutations, the Mutational Signatures Framework [2] that is based on NMF technique, additionally provides estimates of the number of mutations generated by each discovered signature. When a tumor sample is small or one wants to decompose a patient’s mutation profile into already known signatures then the newly developed tool – deconstructSigs (dS), can be used [14]. As both methods are based on heuristics they do not always show consistent levels of signature exposures. In addition, the decomposition problem into known signatures can be solved optimally using existing optimization techniques based on quadratic programming or simulated annealing (see Materials and Methods) providing theoretically the best decomposition solution. We ran all four methods and compared them on the biggest available whole-genome dataset that contains somatic mutations from 560 breast cancer patients [16] (Figure 1).

**Fig. 1.**
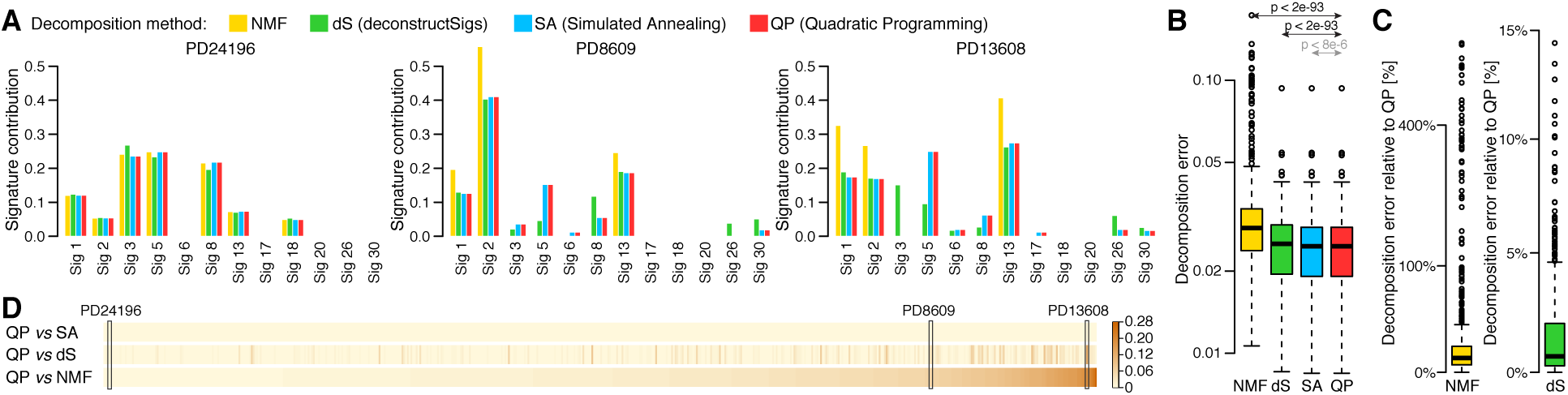
Comparison of four decomposition methods. Decomposition of patient’s mutational profile into 12 mutational signatures known to be present in breast cancers was inferred using four different methods for each of 560 patients. (**A**) Three examples comparing four methods shown as barplots. (**B**) Distribution of decomposition errors (the root sum-squared error between observed and inferred mutational profile) compared between four method across 560 breast cancer patients. Statistical significance of difference between decomposition errors of QP and three other methods is shown (paired Wilcoxon test); Cohen’s d effect size of difference between decomposition errors of QP and SA is negligible. (**C**) Decomposition errors for NMF and dS relative to QP errors shown in the log scale (y axis). (**D**) Comparison of cosine distance between signature contributions inferred using QP and three other methods shown for all patients. Patients were sorted based on their distance between QP and NMF solutions. Three patients selected as the examples in (**A**) are marked.

Results across all methods are often very similar, as for example for patient PD24196 in Figure 1A where the methods show similar contributions of all signatures. However, in many cases their results are significantly different like for patients PD8609 and PD13608 in Figure 1A, where contributions of some signatures vary greatly between methods. QP and SA show very consistent results. NMF identifies the strongest mutational signatures (1, 2, and 13) in each sample, so their exposures are over-estimated as compared to other methods, and it ignores signatures with little contribution. However, signature 5, estimated by QP and SA to have a high contribution (about 0.15 and 0.25 in both examples), is missed. dS usually shows similar results to QP and SA, but for some signatures it differs significantly. For example, in patient PD13608, the signature 3 is detected by dS with high contribution of 0.15 and absent in results of other methods, while the signature 5 is only at 0.1 even though QP and SA shows strong contribution of 0.25. The decomposition errors (values of the objective function) across all 560 patients for four methods are shown in Figure 1B. Although QP finds slightly better solutions than SA in terms of optimal decomposition error in majority of patients (75%), the Cohen’s d effect size of the difference between two methods is negligible so we can assume that they perform equally well. Moreover, QP and SA have always lower decomposition error than two other methods across all 560 patients, and they significantly outperform them in terms of both p-value (<2e-93; paired Wilcoxon test) and Cohen’s d effect size (>0.4). NMF has on average the largest decomposition error that is much higher (by up to 750% and mean of 40%) relative to QP and SA as shown in Figure 1C. However, the main objective of NMF is to discover unknown signatures from patient’s catalogs of mutations, not to find optimal decomposition into predefined signatures. That is why the method focuses on the most prominent signatures in each patient, so the lower performance in terms of optimal error is not surprising. While it is not striking in Figure 1B, the decomposition error of dS is higher by up to 15% (mean 1.6%) relative to QP and SA (Figure 1C). As it is shown in two examples in Figure 1A, even such small differences in decomposition error can lead to significant discrepancy in inferred contributions or even overall signature presence. To show the extent of differences between methods we computed the pairwise cosine distance between exposures inferred by QP versus NMF, dS and SA for each individual patient (Figure 1D). There is a number of patients for which the distance between solutions is large, but it is not specific to particular patients but rather to compared decomposition methods.

The above results show that for some patients the signature exposures inferred by different methods differ significantly, even when there is no striking difference in decomposition errors. Moreover, it appears that some signatures (e.g. 3 and 5) are more prone to variability than other signatures (e.g. 1, 2 and 13). Even though the method based on QP shows the best performance in terms of smallest decomposition error and it is much faster than other methods, it is still not clear how stable the optimal solution is and which signatures are credible. To answer these questions we analyzed the stability of the optimal solution in terms of variability in both input data and suboptimal solutions. This will help us to understand the origin of the observed discrepancies between the methods.

### 3.2 Confidence and stability analysis of signature contributions

The observed mutation frequencies in real biological samples may be contaminated by noise and the optimal solutions inferred based on such data do not always have meaningful and direct biological interpretation. In order to measure the confidence in the estimation of the exposure intensities, we applied the bootstrap technique. Thus we perturbed the original mutational catalog of each patient separately by randomized re-sampling with replacement and we estimated signature contributions in each bootstrap sample using QP method. Figure 2A shows the distribution of exposure estimates in perturbed input catalogs for the same three examples shown in Figure 1A. The bootstrap estimates are, as expected, distributed around the optimal QP solutions in the original data and their median values are reasonably close to the original contributions. However, the variability of the bootstrap solutions for some signatures and patients vary significantly from the original QP exposures. This variability seems to be related to specific signatures and not depended on patients or exposure levels. For example, the contributions of signature 3 differ between the example patients in Figure 2A from 0 to over 0.2, but the variability of bootstrap solutions seems to be high in all three cases. On the other hand, variability of signature 13 contributions in bootstrap samples is small across all examples and also independent of exposure levels. To assess the signature stability more broadly we investigated divergence of their bootstrap contributions from the optimal QP solution in terms of the mean squared error for each patient separately (Figure 2B). Signatures 3, 5 and 8 show large instability in majority of patients; moderate instability can be also observed for signatures 1 and 30. On the contrary, signatures 2 and 13 are stable across all patients. Some of the other signatures (e.g. signature 17) seems to be stable as well, but they are rather infrequent signatures with median contrinution below 0.01.

**Fig. 2.**
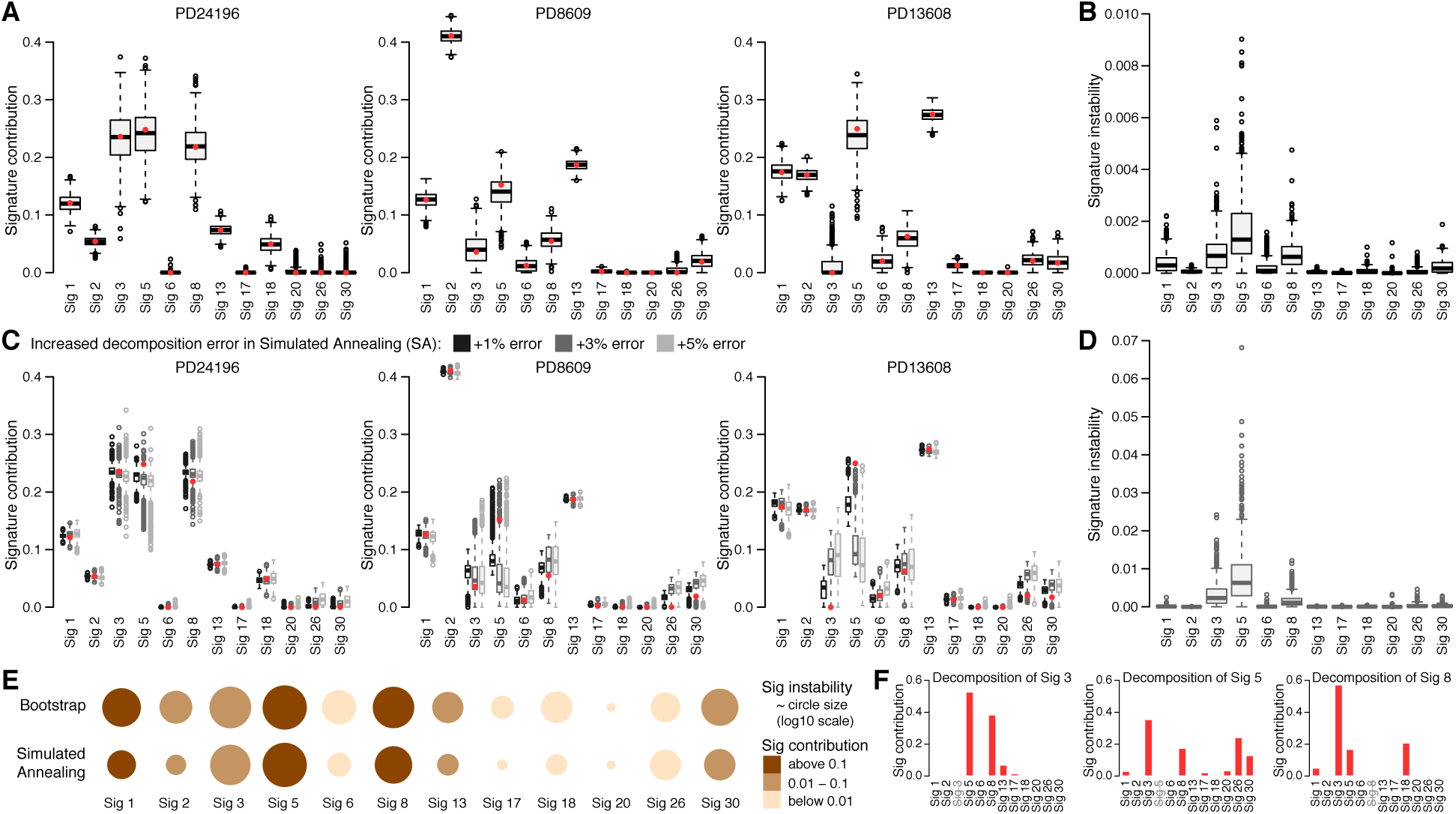
Stability of signature contributions. (**A**) Distribution of signature contributions in 1000 bootstrap samples for the three selected patients (same as in Figure 1A). Contributions in the original samples inferred by QP are shown as red dots. (**B**) Comparison of signature contribution instability between all signatures measured as the mean squared error between contributions in 1000 bootstrap samples and contributions in the original samples for each patient. Corresponding results for suboptimal solutions based on simulated annealing trials with increased decomposition error are shown in (**C**) and (**D**), respectively. Three error threshold increases (+1%, +3% and +5%) relative to the optimal error from QP methods are compared (**C**), but only results of “+3% error” are further presented. (**E**) Summary of stability analysis based on bootstrap (top) and simulated annealing (bottom). The size of each circle represents median signature contribution instability (log10 scale) over all patients for each signature separately; the sizes were normalized by the most unstable signature. The color of each circle indicates median signature contribution. (**F**) Decomposition of a mutational signature (3, 5 or 8) into 11 remaining signatures using QP method.

The COSMIC signatures were derived from a large but limited dataset containing different cancer types using a heuristic method based on NMF, so although the signatures are linearly independent they should not be assumed to be "orthogonal" and some weak dependencies between them might exist. In such case, different linear combinations of signatures can lead to equivalent approximations of the optimal decomposition, i.e. approximations with the same decomposition error in the close proximity of the optimal solution. Then, our confidence in the applicability of the optimal decomposition will be weakened. To check whether this problem applies to all signatures or is caused by only some of them we randomly sampled the solution space of suboptimal decompositions for all patients using simulated annealing and stopping the simulations whenever the decomposition error was close to the error of optimal QP decomposition. We performed three runs of simulations with increasing error of suboptimal solutions – optimal error times 1.01, 1.03, and 1.05, respectively. The results for our three exemplary patients are presented in Figure 2C. Summary of signature stability analysis in terms of MSE for 3% error increase is presented in Figure 2D, but results for all three error levels are consistent. And again, we can observe that contributions of some signatures in suboptimal decompositions, like signatures 2 and 13, are close to the contributions inferred by QP method and that they are stable independently of patient, exposure level and increased error level. This time contributions of signatures 1 and 30 seem to be more stable than in bootstrap analysis. However, the contributions of signatures 3, 5 and 8 in suboptimal decompositions still vary substantially across all patients and in addition they diverge from the optimal QP exposures (see for example patients PD8609 and PD13608). This divergence increases with the increased error of suboptimal decompositions and can quickly lead to over- or underestimation of contributions of some signatures like for example in patient PD13608 for signatures 3 and 5, respectively. This explains why the contributions of signatures 3 and 5 inferred by dS for this patient differed significantly from the contributions inferred by QP (Figure 1A), namely dS found a suboptimal solution with decomposition error higher than in the optimal solution found by QP.

Presented applications of both bootstrap and simulated annealing approaches provide different but complementary views on stability of contributions of mutational signatures inferred from patient’s mutational profile. Contributions of signatures 3, 5 and 8 are unstable from both perspectives (Figure 2E) and their variability seems to be interrelated especially in simulated annealing analysis (Figure 2C). To check whether any signature can be easily replaced by a combination of other signatures we decomposed each signatures into remaining signatures using QP approach. The only signatures that can be decomposed with relatively small error (within decomposition errors of real BRCA patients) are signatures 3, 5 and 8 (Figure 2F). Each of these signatures requires considerable presence of the two other signatures plus limited contribution from some of the remaining signatures. This helps to explain the exposure instability of these three signatures. Moreover, both analyses show that contributions of signatures 2 and 13, both related to the activity of AID/APOBEC, are stable across all patients and that they cannot be replaced by combination of other signatures. This suggest that these two signatures are distinctively defined and well separated from other signatures. Similar properties apply, to some extent, to signatures 1 and 30 that suggests why their instability is reduced in simulated annealing versus bootstrap analysis.

In both analyses, we ran 1000 simulations and computed relevant decompositions for each patient, like for example for three patients in Figures 2A,C. Based on these computations, one can estimate confidence intervals for each signature contribution in each patient to better assess which signatures are present in patient’s mutational profile and what their contribution levels are. Alternatively, one can ask what is the probability that a patient’s exposure to a signature is above a certain level. We counted the number of patients with different exposure levels of all BRCA signatures (Table 1). As we can observe, huge majority of patients (above 85%) is exposed to signatures 1, 5 and 8 at least at a minimal exposure level of 0.01 (with p-value 0.01), and these signatures are relatively abundant in a number of patients. Other signatures like 6, 17, 20 and 26 are only present in limited number of patients and only few patients are exposed to these signatures at levels above 0.15. Such analysis allows to assess which signatures are present in a single sample at a minimal exposure level, so further analysis could be focused on essential signatures only and signatures with little contribution could be disregarded from analysis of a particular patient.

**Table 1.** The number of breast cancer patients exposed to each of 12 mutational signatures (columns) at different minimal contribution levels (rows) with p-value of 0.01 as assessed by bootstrap analysis of the perturbed patient’s mutational catalogs.

| Exposure threshold | Signatures |  |  |  |  |  |  |  |  |  |  |  |
| --- | --- | --- | --- | --- | --- | --- | --- | --- | --- | --- | --- | --- |
|  | 1 | 2 | 3 | 5 | 6 | 8 | 13 | 17 | 18 | 20 | 26 | 30 |
| >0.01 | 512 | 366 | 220 | 505 | 16 | 479 | 402 | 30 | 107 | 5 | 25 | 115 |
| >0.05 | 411 | 173 | 187 | 459 | 7 | 374 | 218 | 8 | 41 | 4 | 11 | 20 |
| >0.10 | 342 | 110 | 164 | 368 | 6 | 225 | 118 | 4 | 16 | 3 | 8 | 2 |
| >0.15 | 258 | 71 | 144 | 301 | 5 | 121 | 78 | 3 | 6 | 1 | 7 | 1 |
| >0.20 | 159 | 50 | 128 | 219 | 4 | 65 | 60 | 2 | 3 | 1 | 7 | 1 |
| >0.25 | 75 | 35 | 118 | 144 | 1 | 27 | 45 | 1 | 1 | 1 | 5 | 1 |

### 3.3 Re-analysis of mutational signatures presence in breast cancers

Nik-Zainal *et al.* [16] analyzed whole-genome sequences of 560 breast cancers and extracted 12 base substitution mutational signatures from patient’s catalogs of somatic mutations using their own framework based on NMF [2]. The method focuses only on the essential mutational signatures that do contribute large numbers of mutations to the inferred mutational profile of each sample. Using the tools we presented in our paper we re-analyzed the data using all 30 COSMIC mutational signatures and determined which other signatures, beyond the 12 known signatures, are present in breast cancers with high confidence. Figure 3A shows distribution of sample exposures to 30 signatures over all patients as inferred by QP method. Three additional signatures 9, 12 and 16, formerly detected in other cancer types, are likely to be present in a number of patients as their contribution levels are comparable to or even exceed contributions of some of the 12 signatures. The results of bootstrap simulations confirmed these observations, see Table 2; and simulated annealing analysis of suboptimal solutions showed consistent results (not shown). As we can observe, each of these novel signatures is present in at least few patients with high contribution above 0.15 and empirical p-value of 0.01; the same is true for the 12 known breast cancer signatures. The remaining signatures do not show the same presence in breast cancer patients, so they were excluded from the further analysis and we recomputed the data decomposition into 15 signatures – 12 known plus 3 novel, for all patients.

**Fig. 3.**
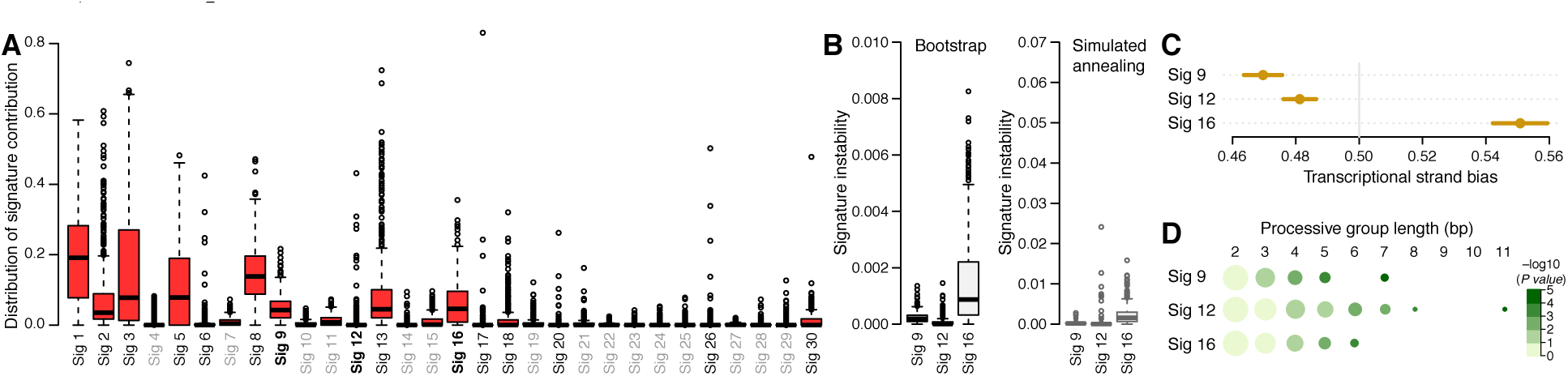
(**A**) Distribution of contributions of 30 COSMIC signatures over 560 breast cancer samples inferred using QP method. Three signatures (9, 12 and 16) suggested to be novel in breast cancers are marked bold; signatures not present in breast cancer are gray. The signatures 9, 12 and 16 were evaluated in terms of their contribution stability in the bootstrap and simulated annealing analyses (**B**) and genomic features known to be exhibited by some signatures such as the transcriptional strand bias (**C**) and length of processive groups (**D**). Observed transcriptional strand bias is shown as a circle with 95% confidence intervals against expected bias of 0.5. Processive groups of different lengths (columns) for each signature (rows) are represented as circles whose size corresponds to the number of groups (log10 scale) and color to the p-value of detecting a processive group of a defined length (-log10 scale). The numbers of groups were normalized by the number of groups of length 2.

**Table 2.** The number of breast cancer patients exposed to each of 30 COSMIC mutational signatures (columns) at different minimal contribution levels (rows) with p-value of 0.01 as assessed by bootstrap analysis of the perturbed patient’s mutational catalogs. The 12 known breast cancer signatures are highlighted in gray and the three signatures (9, 12 and 16) suggested to also be present in breast cancers – in blue.

| Exposure threshold | Signatures |  |  |  |  |  |  |  |  |  |  |  |  |  |  |  |  |  |  |  |  |  |  |  |  |  |  |  |  |  |
| --- | --- | --- | --- | --- | --- | --- | --- | --- | --- | --- | --- | --- | --- | --- | --- | --- | --- | --- | --- | --- | --- | --- | --- | --- | --- | --- | --- | --- | --- | --- |
|  | 1 | 2 | 3 | 4 | 5 | 6 | 7 | 8 | 9 | 10 | 11 | 12 | 13 | 14 | 15 | 16 | 17 | 18 | 19 | 20 | 21 | 22 | 23 | 24 | 25 | 26 | 27 | 28 | 29 | 30 |
| >0.01 | 523 | 335 | 250 | 7 | 91 | 8 | 10 | 434 | 246 | 13 | 18 | 15 | 386 | 5 | 12 | 117 | 9 | 66 | 3 | 5 | 6 | 0 | 0 | 0 | 8 | 7 | 0 | 4 | 2 | 8 |
| >0.05 | 423 | 165 | 210 | 1 | 62 | 6 | 0 | 312 | 65 | 0 | 0 | 9 | 210 | 2 | 1 | 32 | 4 | 25 | 0 | 3 | 3 | 0 | 0 | 0 | 0 | 5 | 0 | 1 | 0 | 1 |
| >0.10 | 347 | 104 | 175 | 0 | 33 | 6 | 0 | 174 | 12 | 0 | 0 | 8 | 117 | 0 | 0 | 8 | 3 | 9 | 0 | 2 | 1 | 0 | 0 | 0 | 0 | 4 | 0 | 0 | 0 | 1 |
| >0.15 | 265 | 69 | 154 | 0 | 12 | 4 | 0 | 91 | 2 | 0 | 0 | 7 | 78 | 0 | 0 | 2 | 3 | 5 | 0 | 1 | 0 | 0 | 0 | 0 | 0 | 3 | 0 | 0 | 0 | 1 |
| >0.20 | 177 | 49 | 136 | 0 | 4 | 2 | 0 | 36 | 0 | 0 | 0 | 4 | 61 | 0 | 0 | 1 | 2 | 1 | 0 | 1 | 0 | 0 | 0 | 0 | 0 | 2 | 0 | 0 | 0 | 1 |
| >0.25 | 96 | 35 | 120 | 0 | 1 | 2 | 0 | 16 | 0 | 0 | 0 | 3 | 45 | 0 | 0 | 0 | 1 | 1 | 0 | 0 | 0 | 0 | 0 | 0 | 0 | 2 | 0 | 0 | 0 | 1 |

To further evaluate the presence of novel signatures 9, 12 and 16 in breast cancers we assessed their stability and we tested if these signatures exhibit some known transcriptional features previously shown for some of the 12 signatures by Morganella et al. [17]. The novel signatures show similar range of stability as the 12 signatures, from stable signature 12 to unstable signature 16 (Figure 3B; compare with Figure 2B,D). Transcriptional strand bias of the novel signatures is highly significant (p-value < e-12) and much stronger than for any other signature (Figure 3C). Signatures 9 and 12 show bias towards the non-transcribed strand, while signature 16 shows extremely strong bias towards the transcribed strand, mostly with T>C mutations at ATN context. The bias of signatures 12 and 16 was formerly observed in other cancer types, while the bias of signature 9 was not observed before and it seems to be associated with the function of DNA polymerase *η* that plays an essential role in replicating damaged DNA. Some mutational processes cause long stretches of successive mutations occurring on the same DNA strand, called processive groups. We found significantly long processive groups containing at least 5-6 mutations caused by the novel signatures, especially for signature 12 that had the longest processive group of 11 substitutions (Figure 3D). These results shows that the novel signatures, 9, 12 and 16, carry genomic features previously observed for breast cancer signatures or signatures present in other cancer types and independently support the outcome of our tools for decomposition of patient’s mutational profile into predefined mutation signatures with confidence.

## 4 Conclusions

With the steadily decreasing sequencing cost, we can now catalog cancer somatic mutations for ever increasing number of individual patients. It is important to be able to use this information optimally. In particular, key information that we can obtain from such mutation data are the mutagenic processes shaping the mutation catalog of each individual patient. Such information can point to specific defects in the replication mechanism, enzymatic activities, etc. Until now methods that decomposed patient’s mutational catalog into mutations associated with specific signatures made no attempt to assess confidence in such decompositions. We showed that the results provided by existing methods can differ significantly not only in estimates of exposure levels to each mutagenic process but even in assessing whether or not a given signature is actually present. In particular, we provided statistical evidence and biological support for the presence of three additional signatures in breast cancers. However, many of the differences are not merely a reflection of differences in the accuracy of decomposition methods, but are at least in part related to the inherent instability of some signatures.

In this work, we used two complementary approaches to rigorously assess confidence and stability of the resulting decomposition. Our analysis showed that some mutagenic signatures, such as the signatures related to the APOBEC activity, are very stable but some are not. Especially noisy are signatures 3, 5 and 8. This instability can be explained, in part, by the fact that these signatures can be decomposed into linear combination of other signatures with a very small error. This problem can only deepen with the increasing number of novel mutational signatures that are being discovered thanks to steadily increasing cancer datasets. Our results emphasize the importance of analyzing confidence and stability of inferred signature contributions from the perspective of input data perturbation and approximate suboptimal solutions and the evaluation methods and software developed for this study can aid such analyses.

## Supplementary Materials

### Quadratic Programming (QP)

Taking advantage of the COSMIC mutational signature matrix *P*, as columns of *P* are linearly independent so *P ^T^ P* is positive definite, our minimization problem is equivalent to optimization of a strictly convex quadratic problem. Dual method [15] is an excellent technique to get an efficient and numerically stable solutions to this kind of quadratic problems by utilizing the Cholesky and QR factorizations and updating procedures. Our objective function can be re-written as:

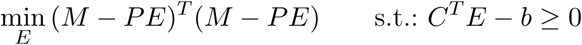
where
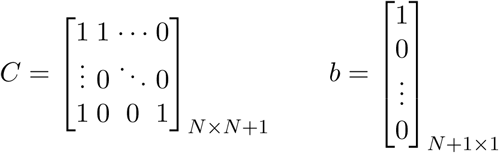
and the first constraint here is an equality constraint, and all further are inequality constraints. And further we can re-write the problem in the form of problem solved by the dual method:
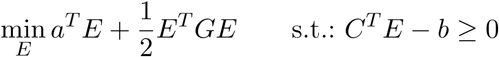
where *a*^*T*^ = *M ^T^ P*, *G* = *P ^T^ P*. Note that *G* is positive definite for the COSMIC matrix *P*.

By calling function solve. QP from the R package *quadprog*, our optimization problem is solved very fast in less than 30 iterations on average. Similar approach based on QP was also suggested by Lynch [18].

### Simulated annealing (SA)

One of popular optimization algorithms is simulated annealing. We used the generalized simulated annealing (GSA) approach [19] implemented in the R package *GenSA* [20]. GSA relies on Tsallis statistics [19], which is a generalization of the Boltzmann-Gibbs statistics used in classical SA. GSA is able to accurately locate the absolute minimum of a given function and convergence is reached much more rapidly than in classical SA.

## References

[1] S. Nik-Zainal et al. “Mutational processes molding the genomes of 21 breast cancers”. In: Cell 149.5 (2012), pp. 979–93.

[2] L. B. Alexandrov et al. “Deciphering signatures of mutational processes operative in human cancer”. In: Cell Rep 3.1 (2013), pp. 246–59.

[3] L. B. Alexandrov et al. “Signatures of mutational processes in human cancer”. In: Nature 500.7463 (2013), pp. 415–21.

[4] T. Helleday, S. Eshtad, and S. Nik-Zainal. “Mechanisms underlying mutational signatures in human cancers”. In: Nat Rev Genet 15.9 (2014), pp. 585–98.

[5] L. B. Alexandrov and M. R. Stratton. “Mutational signatures: the patterns of somatic mutations hidden in cancer genomes”. In: Curr Opin Genet Dev 24 (2014), pp. 52–60.

[6] A. Fischer, C. J. Illingworth, P. J. Campbell, and V. Mustonen,. “EMu: probabilistic inference of mutational processes and their localization in the cancer genome”. In: Genome Biol 14.4 (2013), R39.

[7] S. A. Roberts et al. “An APOBEC cytidine deaminase mutagenesis pattern is widespread in human cancers”. In: Nat Genet 45.9 (2013), pp. 970–6.

[8] N. Kanu et al. “DNA replication stress mediates APOBEC3 family mutagenesis in breast cancer”. In: Genome Biol 17.1 (2016), p. 185.

[9] N. J. Haradhvala et al. “Mutational Strand Asymmetries in Cancer Genomes Reveal Mechanisms of DNA Damage and Repair”. In: Cell 164.3 (2016), pp. 538–49.

[10] S. Henderson et al. “APOBEC-mediated cytosine deamination links PIK3CA helical domain mutations to human papillomavirus-driven tumor development”. In: Cell Rep 7.6 (2014), pp. 1833–41.

[11] M. B. Burns et al. “APOBEC3B is an enzymatic source of mutation in breast cancer”. In: Nature 494.7437 (2013), pp. 366–70.

[12] Y. A. Kim, S. Madan, and T. M. Przytycka. “WeSME: uncovering mutual exclusivity of cancer drivers and beyond”. In: Bioinformatics 33.6 (2017), pp. 814–821.

[13] L. B. Alexandrov et al. “Mutational signatures associated with tobacco smoking in human cancer”. In: Science 354.6312 (2016), pp. 618–622.

[14] R. Rosenthal et al. “DeconstructSigs: delineating mutational processes in single tumors distinguishes DNA repair deficiencies and patterns of carcinoma evolution”. In: Genome Biol 17 (2016), p. 31.

[15] D. Goldfarb and A. Idnani,. “A numerically stable dual method for solving strictly convex quadratic programs”. In: Mathematical Programming 27.1 (1983), pp. 1–33.

[16] S. Nik-Zainal et al. “Landscape of somatic mutations in 560 breast cancer whole-genome sequences”. In: Nature 534.7605 (2016), pp. 47–54.

[17] S. Morganella et al. “The topography of mutational processes in breast cancer genomes”. In: Nat Commun 7 (2016), p. 11383.

[18] A. G. Lynch. Decomposition of mutational context signatures using quadratic programming methods [version 1; referees: 1 approved, 1 approved with reservations]. Vol. 5. 2016.

[19] C. Tsallis and D. A. Stariolo. “Generalized simulated annealing”. In: Physica A, 233 (1996), pp. 395–406.

[20] Y. Xiang, S. Gubian, B. Suomela, and J. Hoeng,. “Generalized Simulated Annealing for Efficient Global Optimization: the GenSA Package for R.” In: The R Journal Volume 5/1, June 2013 (2013).

